# *In flagranti* - Functional morphology of copulatory organs of odontopygid millipedes (Diplopoda: Juliformia: Spirostreptida)

**DOI:** 10.1101/2022.11.22.517485

**Authors:** Justus Brandt, Hans Simon Reip, Benjamin Naumann

## Abstract

The copulatory organs of many animal groups exhibit a high degree of morphological complexity and diversity that is thought to have evolved on the basis of different selective mechanisms including lock-and-key mechanism, pleiotropy, sperm competition, internal courtship and female choice. Identifying the effects of these different selective mechanisms on copulatory organs one of the central topics of the study of sexual selection. To tackle this challenge, knowledge of the functions of all parts of the copulatory organs is indispensable. Here we study the functional morphology of the gonopods (male copulatory organs) and vulvae (female copulatory organs) in the odontopygid millipede *Spinotarsus* (Diplopoda, Spirostreptida, Odontopygidae). While the vulvae of female odontopygids are rather simple, male gonopods are complex, walking leg-derived copulatory organs that exhibit many movable sub-parts. Using μCT-based 3D reconstruction, confocal laser scanning microscopy and mating observations we revise the functional morphology of odontopygid gonopods, propose biological roles and evaluate the possible involvement of different selective mechanisms underlying their evolution.

## INTRODUCTION

The copulatory organs of many animal groups exhibit a high degree of morphological complexity and diversity that cannot be explained solely by selection for their general function to transfer or receive sperm (Eberhard, 1985; Tadler, 1993). Additionally, sexual selection is thought to play a major role in the evolution of morphologically complex copulatory organs (Eberhard, 1985, 1993). Within this frame, theories based on different selective mechanisms such as the lock-and-key mechanism, pleiotropy, sperm competition, internal courtship and female choice have been proposed to explain the rapid divergent evolution observed especially in male copulatory organs (Eberhard, 1985, 1993). These theories impose the presence of intrasexual as well as intersexual competition for the “best-fitting”, most attractive and most competitive male copulatory organs. Entangling the effects of the different selective mechanisms involved in the evolution of male copulatory organ morphology remains one of the central challenges to the study of sexual selection (Telford and Dangerfield, 1996). To tackle this challenge, knowledge of the functions of all (sub-) parts of copulatory organs during mating is indispensable. However, most of the proposed functions are based on detailed morphological descriptions of isolated male and female copulatory organs. Consequently, studies examining interacting male and female copulatory organs are crucial to test if *functions* proposed based on isolated specimens carry out *biological roles* (Bock and von Wahlert, 1965).

The copulatory organs of male helminthomorph diplopods are called gonopods and exhibit a high degree of morphological complexity and disparity used for taxonomic species discrimination (Attems, 1914; Attems, 1950; Kraus, 1966; Minelli, 2015). There is still a debate about the selective mechanisms underlying the evolution of this disparity. Verhoeff (1926-32) favoured a lock-and-key mechanism, Kraus (1966) proposed pleiotropy, and Tadler (1993, 1996), Barnett and Telford (1997), Barnett (1997), Frederiksen and Enghoff (2015) argued for sperm competition, internal courtship and female choice (Figure 1). Recent studies using three-dimensional (3D) models based on micro-computed tomography (μCT) retrieved the idea of a lock-and-key mechanisms in a paradoxomatid helminthomorph (Wojcieszek et al., 2012) or proposed a combination of lock-and-key and female choice in a polydesmid helminthomorph (Zahnle et al., 2020).

**Figure 1.**
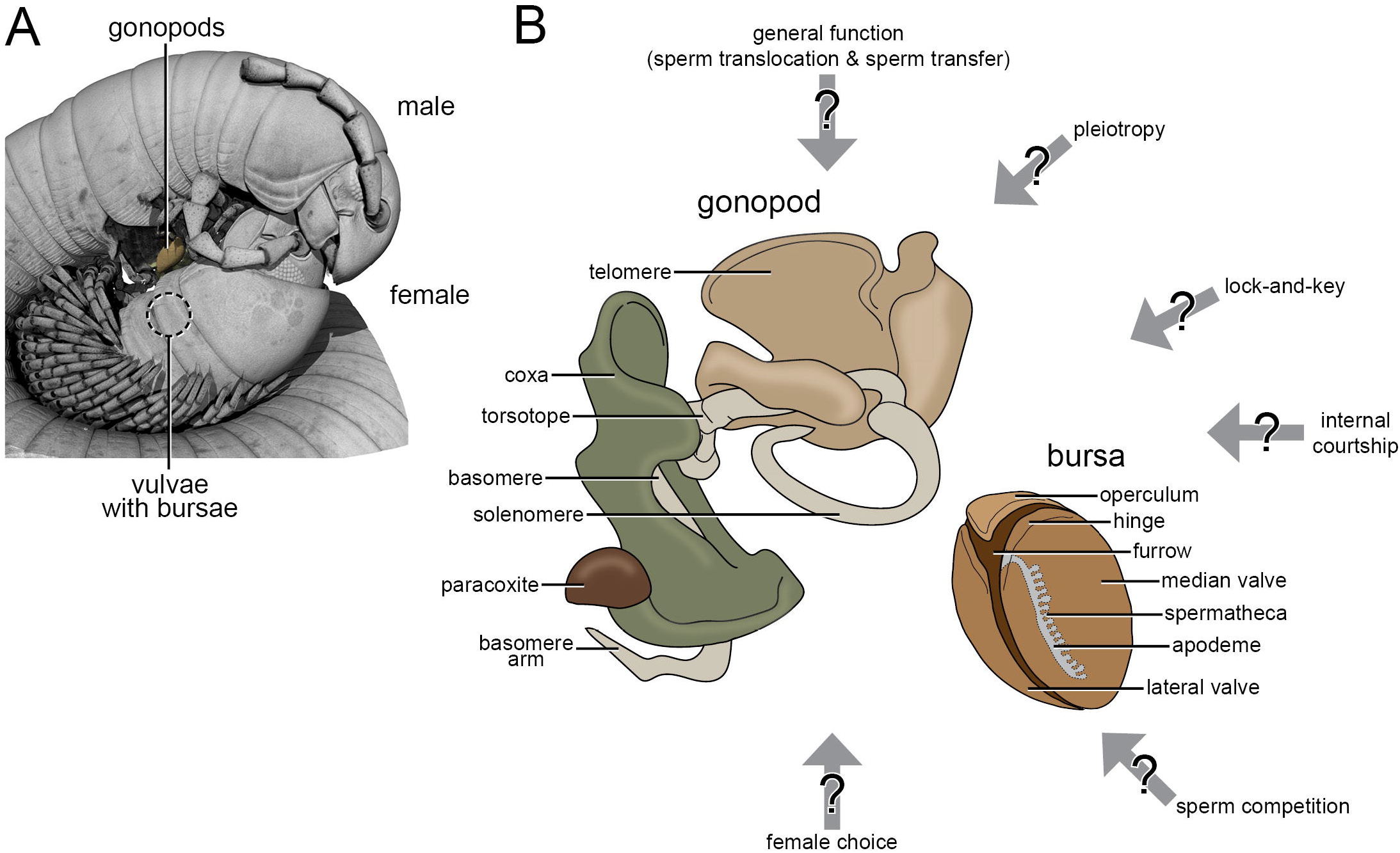
A, volume rendering of a copulating pair of S. *glabrus*. B, schematic illustration of a male gonopod and a female bursa of *S. glabrus*. Grey arrows indicate the possible influence of different selective mechanisms on the morphology of both copulatory structures.

In this study, we investigate the functional morphology of male gonopods and female vulvae in copulated pairs of *Spinotarsus glabrus* Kraus, 1960 (Juliformia, Spirostreptida, Odontopygidae). Copulating pairs of other Odontopygidae have been studied previously based on dissections (Barnett, 1997; Barnett and Telford, 1996; Kraus, 1966; Kraus, 1968). Using μCT-based 3D reconstructions in combination with confocal laser scanning microscopy we revise the functional morphology of gonopods of the Odontopygidae, propose biological roles and evaluate the possible involvement of different selective mechanisms underlying their evolution.

## MATERIAL AND METHODS

### SPECIMENS

Specimens determined as *S. glabrus* were collected at the Spion Kop Lodge, South Africa (−28.6952, 29.5365) in November 2018. Living specimens were sexed and separated for a few hours. A few uncopulated males and females were fixed and stored in 70% Ethanol. Additionally, some other specimens were transferred in pairs in small plastic containers (15 × 15 × 5 cm) lined with a wet paper tissue for mating observations.

### MATING VIDEO RECORDINGS

Mating of at least 15 mating pairs was observed during a period of two weeks to describe the mating behavior of *S. glabrus*. Recordings of two pairs were prepared using a standard Canon 650D camera and a set of standard LED light panels for illumination. During the complete mating recording session, a “Best of Muddy Waters” album was played loudly in the background. Most pairs started copulating immediately or after a few minutes. After around 5 minutes (pair 1) and 10 minutes (pair 2) copulating pairs were shock-frozen using ice spray for sport injuries (Lifemed) and submerged in -20°C cold 70% ethanol. In total, micro-computed tomography scans of two copulating pairs and one uncopulated female as well as confocal laser scanning microscopy scans of the gonopods of two uncopulated males and bursae of one uncopulated female were prepared for this study.

### CONFOCAL LASER SCANNING MICROSCOPY (CLSM)

Dissected gonopods and vulvae stored in 70% ethanol were further dehydrated via an ethanol series (20 minutes in 90%, 96% and 99% ethanol each), transferred to 100% methanol for 30 minutes and afterwards cleared and stored in BABB (benzyl alcohol/benzyl benzoate; 2/1).

Afterwards, the material was mounted in BABB in between two coverslips and scanned using a Stellaris 8 (Leica) at 430 nm, 488 nm and 566 nm wavelengths taking advantage of the samples auto fluorescent properties.

### MICRO-COMPUTED TOMOGRAPHY (μCT)

Specimens were cut in half to improve penetration of the contrasting solution. Afterwards, specimens were contrasted in 0.5 % iodine-solution in 70% ethanol for several days. Immediately prior to scanning specimens were washed in 80% ethanol (three to five times for 20 minutes each) and mounted within Eppendorf tubes of different sizes (1.5 to 5 ml) using an ethanol-soaked paper tissue to keep samples in place during the scans. Scans were done using an Xradia 410 Versa (Carl Zeiss AG, Germany).

### THREE-DIMENSINAL (3D) RECONSTRUCTION

The 3D reconstruction based on digital image stacks obtained from CLSM and μCT scans were carried out using AMIRA 5.4.5 (Visage Imaging GmbH, Germany). Individual image stacks of segmented structures were exported to VGStudio Max 2.0.5 (Volume Graphics GmbH, Germany) for volume renderings.

### IMAGE PROCESSING

Brightness, contrast and coloration were adjusted using either Photoshop (Adobe Inc., USA) or Fiji (Schindelin et al., 2012).

### TERMINOLOGY

For male gonopods the terminology introduced by Frederiksen and Enghoff (2012) was used. For female vulvae we used the terminology used by Barnett (1997). Muscles are named according to Wilson (2002) as far as possible.

## RESULTS

### MATING BEHAVIOR

Mating was observed in fifteen pairs. In 10 cases, mating was successful and can be divided into a pre-copulatory phase, four copulatory phases and a postcopulatory phase. In five cases, mating was unsuccessful and was terminated by the pair in between copulatory phase I and II (for individual times see Supplementary Material 1). The video recording corresponding to single images in Figure 2 is available as Supplementary Material 2.

**Figure 2.**
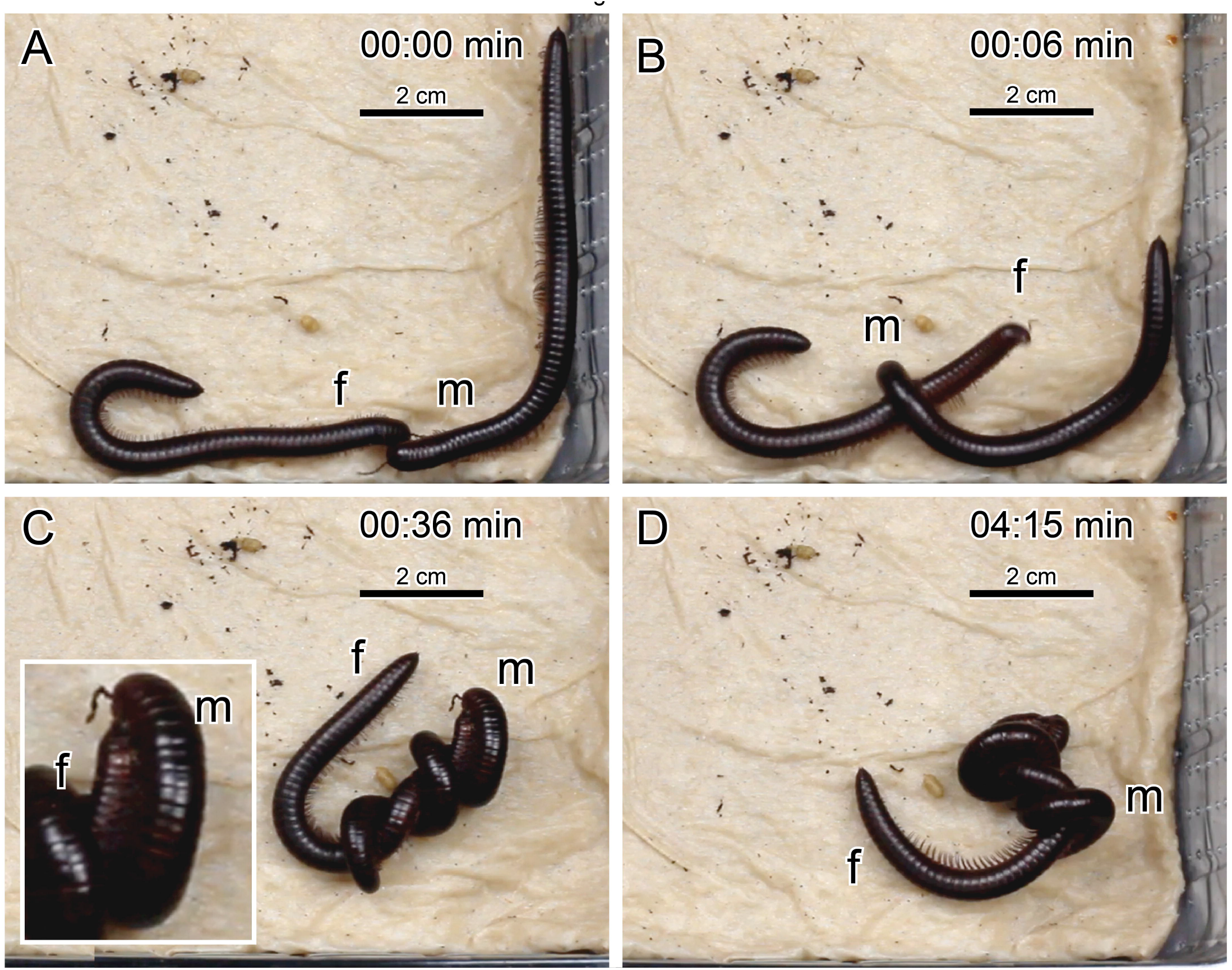
A-D, Photographs of the copulatory sequence of *S. glabrus*. Numbers in the top-right corners indicate the time that passed starting with first tactile contact in A. f, female; m, male.

Pre-copulatory phase – After making contact head-on (Figure 2A), the male starts to quickly and forcefully wrap its body around the female (Figure 2B). About 20 seconds after first contact the entire male is wrapped around the female, reaching up to two full turns (Figure 2C). The male’s head is now located slightly above the female’s, lining up its gonopods in the seventh body ring and the female’s vulvae in the second body ring. Using its antennae, the male continually taps on the female’s head. The female exhibits some escape behavior, twisting and bending its body for approximately 30 seconds to one minute after male wrapping has started. Thereafter, the female stays mostly motionless for the rest of the copulation.

Copulatory phase I – During the first two to three minutes, the male episodically convulses and adjusts its position around the female. Gonopods protrude from the body ring sitting on an inflating hydrostatic gonopodal pillow.

Copulatory phase II – After a suitable position is reached the male moves its right and left gonopod alternately in and out of the female vulvae for around three to five minutes.

Copulatory phase III – The pair stays more or less motionless and gonopods move only infrequently. This phase lasts between approximately 30 to 90 minutes (Figure 2D).

Copulatory phase IV – The male removes its gonopods with an “unlocking”-movement by lowering his anterior body region and moving back from the female.

Postcopulatory phase – Following gonopod removal and the subsequent ending of the copula the male moves away from the female. In some cases, the female resides at the same position cleaning its vulvae with its mouth parts for approximately 20 to 30 minutes.

### UNCOPULATED GONOPOD MORPHOLOGY

Each of the paired gonopods of Odontopygidae can be divided into two main structures, the coxa and the telopodite (Frederiksen and Enghoff, 2012). The uncopulated configuration of a gonopod of *S. glabrus* to which the descriptions correspond is shown in Figure 3.

**Figure 3.**
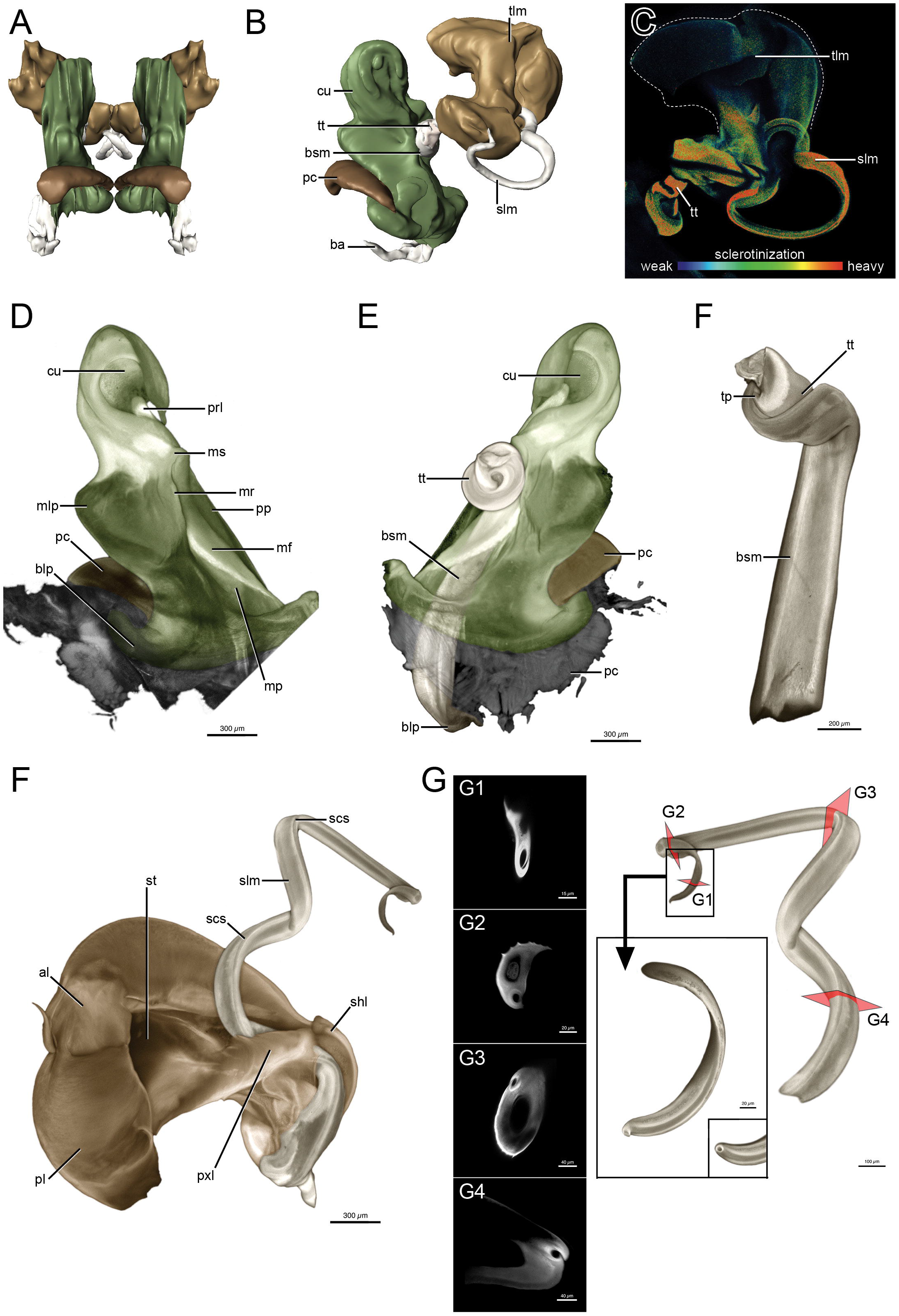
A and B, CLSM-based 3D reconstructions of the uncopulated gonopods of male *S. glabrus*. A, Localization of gonopods in resting position from an anterior view. B, Right gonopod from the lateral view. C, Torsotope, solenomere and telomere od the right gonopod. The colors indicate the degree of sclerotization based on the intensity of autofluorescence at 488 nm. D to G, False-colored maximum intensity projections of CLSM-scans at 488 nm. D, Right coxa and paracoxite from a posterolateral view. E, Left coxa and paracoxite from a posterolateral view. F, Right basomere and torsotope from a medial view. D, Right telomere and solenomere from a posteromedial view. G, Left solenomere from posteromedial view. Red plains indicate transverse sections through the solenomere (G1-G4) to depict the large solenomere channel and the small solenomere channel. The rectangle shows close ups of the solenomere tip to show the opening of the small channel. al, anterior telomeral lamella; ba, basomere arm; blp, basal lateral coxal process; bsm, basomere; cu, cucullus; mf, metaplical flange; mlp, mesal lateral coxal process; mp, metaplica; mr, metaplical ridge; ms, metaplical shelf; pc, paracoxite; pl, posterior telomeral lemalla; pp, proplical; prl, proplical lobe; pxl, proximal telomeral lobe; scs, surface channel of the solenomere; shl, shallow telomeral lobe; slm, solenomere; st, main telomeral stem; tlm, telomere; tp, torsotope process; tt, torsotope

Coxa – The coxa is a dorsoventrad folded sheet consisting of an anterior proplica and a posterior metaplica with their free margins pointing mesad. The metaplica is larger compared to the proplica. It has a large mesal indentation resulting in a posterior-facing ridge next to the free margin below the metaplical shelf. The distal ends of both, the proplica and the metaplica join into a cucullus. The cucullus curves mesad. Its edge is rounded with a triangular protrusion pointing proximad. In mesal view this protrusion reaches over the proplical lobe, thereby covering it. A paracoxite is present at the base of the coxa. Its concave surface points laterally away from the coxa. A matching depression in the coxal surface allows for close contact between the coxa and the paracoxite.

The telopodite consists of four substructures.

Basomere – The basomere is a long straight rod embedded into the coxal lumen formed by the pro- and metaplica. Proximally, it forms a laterally projecting arm curving around the coxal base. The opening of a large channel opens laterodorsad. This large channel continues from the basomere, through the torsotope ending blind almost at the solenomere tip. Some form of a coagulated substance is present close to the blind ending of this channel. A second, much thinner channel is present within the basomere and located posterior and opens close to the large channel. A second smaller opening is present slightly further up on the basomere. After its opening, the small channel submerges again under the surface of the basomere. Shortly before reaching the torsotope, it opens via an infolding of the solenomere. The walls of this infolding make contact with each other and exhibit a slight overlap. This results in a delicate channel running directly along the surface of the solenomere opening at its tip.

Torsotope – The torsotope emerges from a 90° bend of the basomere and is located outside of the coxal lumen. It rests against the proximal side of the metaplical shelf and exhibits one and a half turns. The torsion begins mesad, with the basomere coiling towards the posterior. Some areas of the torsotope appear heavier sclerotized compared to others (Figure 3C). A triangular torsotope process is present pointing laterally. Following the torsotope the telopodite is reduced in diameter in the posttorsal narrowing. Just after this narrowing the solenomere and telomere emerge.

Solenomere – The solenomere is a whip-like structure. After emerging from the torsotope the solenomere first enters the space between the proximal lobe of the telomere and the shallow lobe of the main telomere stem. The solenomere then bends away from the telomere and continues spiral-like for about two and a half turns. The final twists change its direction so that the tip is oriented towards the solenomere base.

Telomere – The telomere originates from the torsotope alongside the solenomere. It features multiple lamellae. Shortly after the point of emergence just after the posttorsal narrowing the proximal lobe and the shallow lobe enclose the basal area of the solenomere. The distal lamellar part of the telomere is folded in itself and features a cleft on its edge, splitting the lamella in two regions. The posterior lamella is larger than the anterior lamella. Close to the cleft a small spine is present. Overall, the telomere is a weakly sclerotized part, compared to e.g., the solenomere (Figure 3C).

### UNCOPULATED VULVAE MORPHOLOGY

The paired vulva of Odontopygidae consist of a thin membranous vulval sac and a bursa each. The sacs open behind the second leg pair. Each sac houses a more or less sclerotized bursa which is connected to the oviduct via a base-region (Barnett, 1997). The uncopulated configuration of the bursa of *S. glabrus* to which the descriptions correspond is shown in Figure 4.

**Figure 4.**
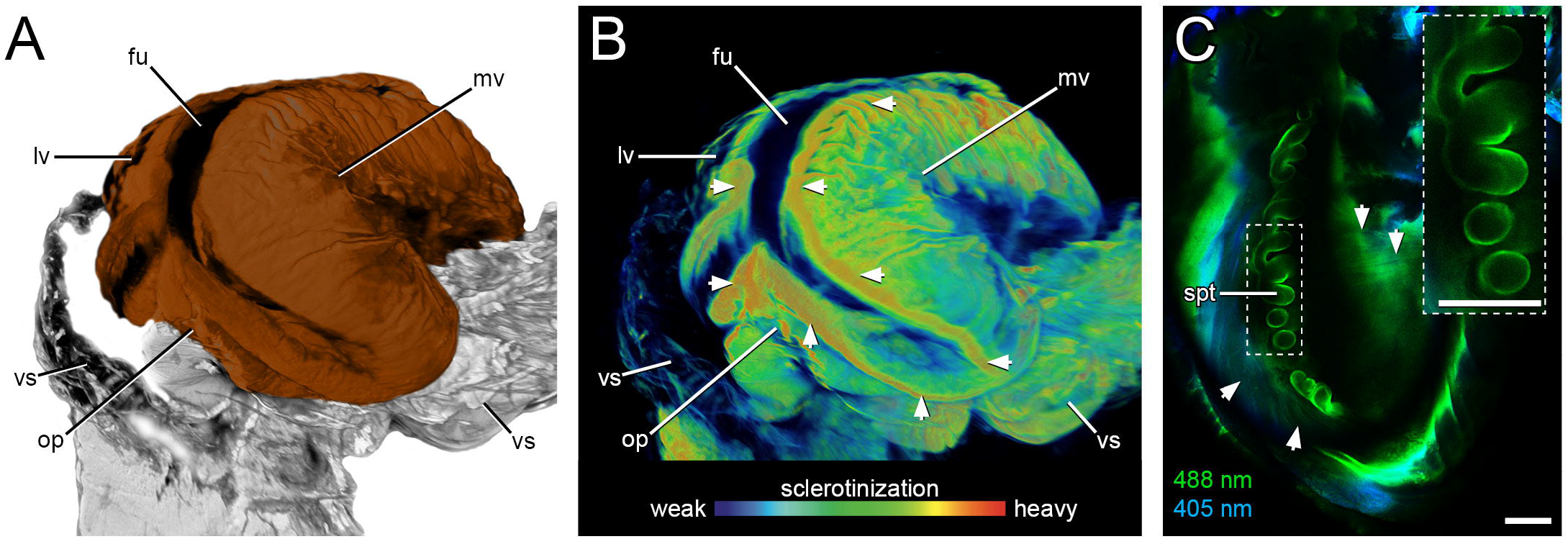
A, False-colored maximum intensity projections of a CLSM-scan of a bursa of female *S. glabrus* at 488 nm. B, The same bursa. The colours indicate the degree of sclerotization based on the intensity of autofluorescence at 488 nm. White arrows indicate the heavily sclerotized hinge. C, CLSM-scan at 405 nm (blue) and 488 nm (green) showing the internal of the bursa. The rectangle in the dashed-lines indicates the position of the close-up of the spermathecae in the right upper corner. White arrows indicate bursal muscles connected to the spermathecae. The scale bar is 100 μm. fu, furrow; lv, lateral valve; mv, median valve; op, operculum; spt, spermathecae; vs, vulval sac.

Bursa – The paired bursae are located deeply within the vulval sac and consist of three weakly sclerotized chitinous plates each. The first - the operculum - covers the opening of the oviduct. It is attached to the other two plates, the median and lateral valve via a more heavily sclerotized hinge that forms a Y-shaped depression in between the three plates (Figure 4B). Anteroventrally, the median and lateral valves are separated by a deep furrow at its dorsal end close to the hinge. It gives rise to the opening of an apodemic tube. This tube connects a series of approximately 20 to 25 small ampullae serving as spermathecae. A broad muscular sheet connects these spermathecae to the bursal wall.

### COPULATED GONOPOD AND VULVAE MORPHOLOGY

During copulation, the male is located above the female with their ventral anterior body regions facing each other. The gonopods protrude from the male’s body via a haemolymph-filled gonopodal pillow. The gonopods from the proximal third of the coxa onwards are located completely inside the female’s vulval sac.

Gonopods (Figure 5 and 6)

**Figure 5.**
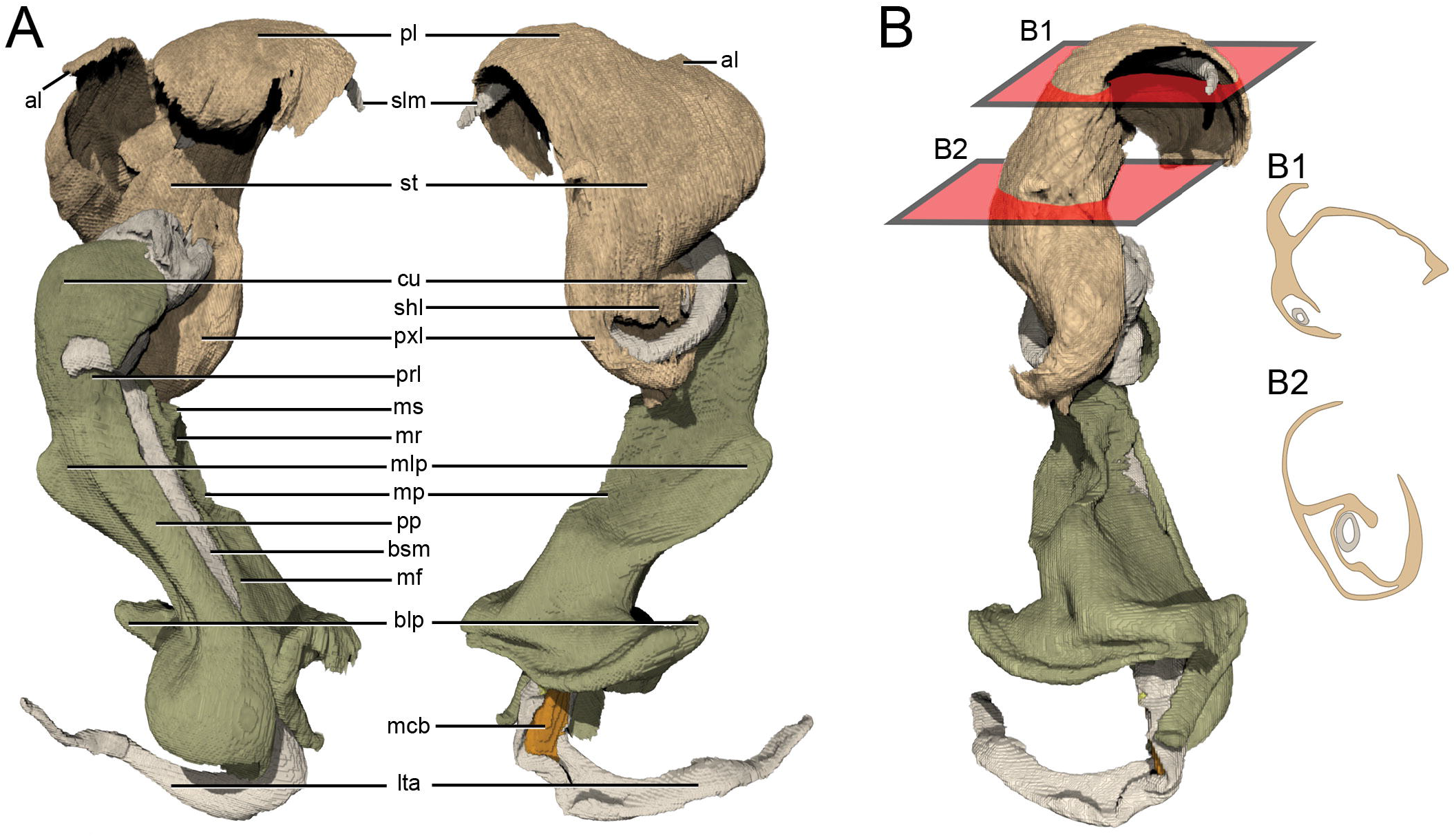
3D reconstructions of the gonopods in copula without the bursae. A, Left gonopod in anteriomesal view (left) and posteriolateral view (right). B, Gonopod from a mesial view. Red planes indicate transverse sections through the telomere and solenomere (B1 and B2) to depict their interaction. al, anterior telomeral lamella; blp, basal lateral coxal process; bsm, basomere; cu, cucullus; lta, lateral telopodite arm; mcb, main channel of basomere; mf, metaplical flange; mlp, mesal lateral coxal process; mp, metaplica, mr, metaplical ridge; ms, metaplical shelf; pl, posterior telomeral lamella, pp, proplica, prl, proplical lobe; pxl, proximal telomeral lobe; scb, small channel of basomere; shl, shallow telomeral lobe; slm, solenomere; st, main telomeral stem.

**Figure 6.**
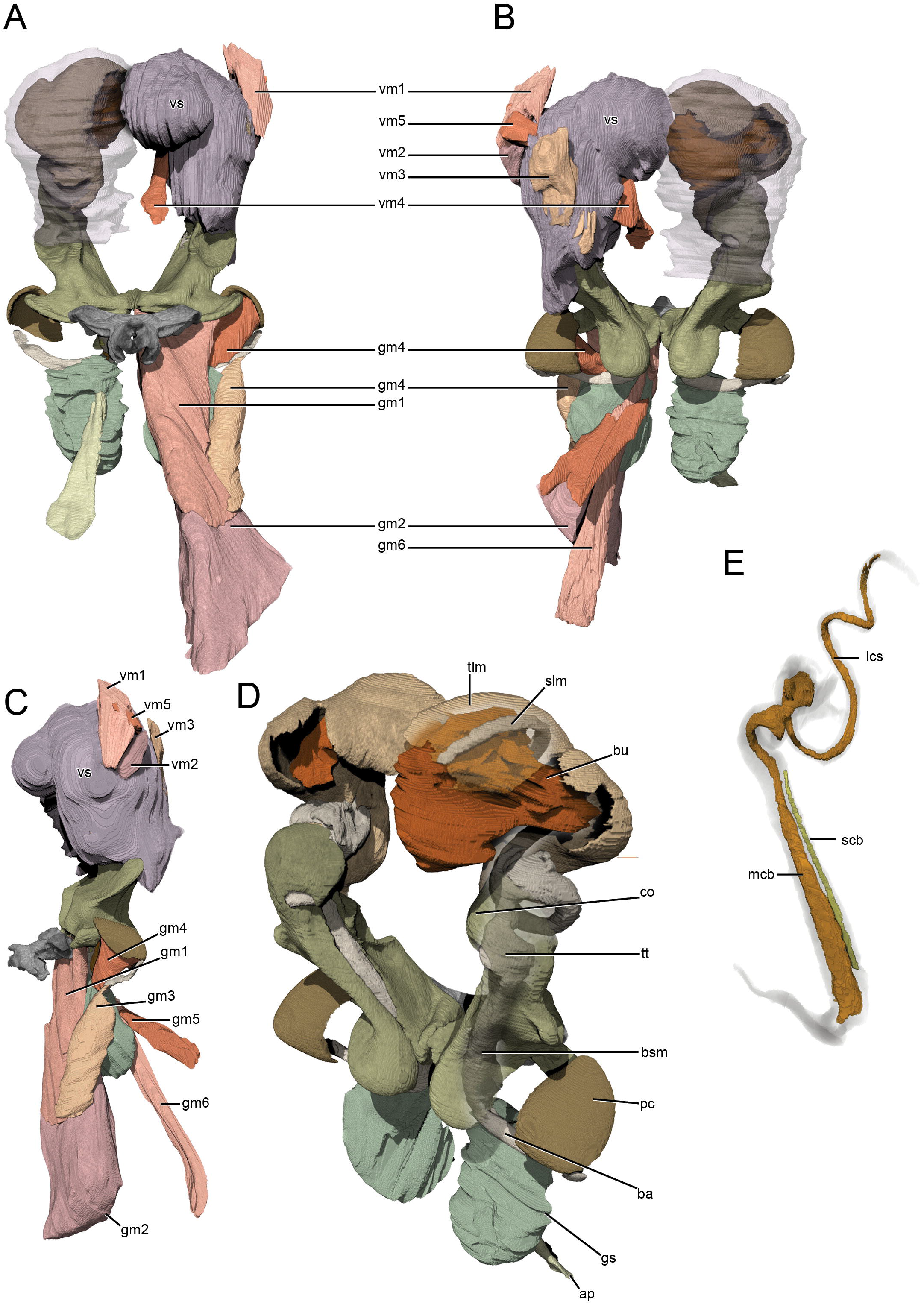
3D reconstructions of the gonopods and vulvae in copula. A-C, Copulatory interface with musculature visible on left gonopod and right vulval sac. Left vulval sac made transparent. A, Posterior view. B, Anterior view. C, Lateral view. D, Copula without musculature and vulval sac. Right telomere and coxa made partially transparent to show telopodite and placement of solenomere above the bursa E, Basomere, torsotope and solenomere made transparent to show the channels inside them. ap, apodeme; ba, basomere arm; bsm, basomere; bu, bursa; co, coxa; gm1, Distal apodeme-coxa muscle; gm2, Apodeme-tergite muscle; gm3, Lateral apodeme-telopodite muscle; gm4, Paracoxite-telopodite muscle; gm5, Ventral t6-gonopodal sac muscle; gm6, Dorsal t6-gonopodal sac muscle; gs, gonopodal sac; lcs, large channel of the solenomere; mcb, main channel of the basomere; pc, paracoxite; scb, small channel of the basomere; slm, solenomere; tlm, telomere; tt, torsotope; vm1, t3-vulval sac ventral muscle; vm2, t3 phragma posterior-vulval sac muscle; vm3, t3 phragma anterior-vulval sac muscle; vm4, t3 sternite-vulval sac muscle; vm5, t3-vulval sac median muscle; vs, vulval sac.

Coxa – Overall little difference can be observed between the copulated and uncopulated coxae. In the copulated state, the cucullus appears to be less curled in. Especially on the posterior side the edge of the cucullus reaches further away from the metaplica compared to the uncopulated coxa. The relationship between coxa and paracoxite is also altered. In comparison to the uncopulated state the lateral edge of the paracoxite appears to have been lowered down proximad and thus projects away laterally with the convex side of the scoop-shape facing distad. The lateral edge of the paracoxite comes close to the tip of the lateral telopodite arm but is not in direct contact with it. The overall form of the paracoxite remains similar.

Major differences can be found between the copulated and uncopulated telopodites and in their interaction with the coxae.

Basomere – The location at which the basomere exits the coxa is shifted towards the distal end of the coxa. The cucullus is wrapped around the torsotope and the proplical lobe enters into the torsotope spiral.

Torsotope – The torsotope introduces much less torsion at only about 180° with the torsotope process bending an additional 90°. Viewed from the anterior side the torsotope introduces almost no directional change to the basomere. From the lateral view the torsotope introduces an approximately 45° posteriad bend.

Solenomere – After emerging from the torsotope the solenomere performs a complicated spiral running first proximad, then mesad and finally curving distad again. The apex of this basally facing loop lies posterior to the basomere and approximately at the height of the torsotope. Contrary to the uncopulated state where the solenomere bends away from the telomere, it is now in close contact to the telomere in copula. The tip of the solenomere rests above the furrow produced by the apodemic tube (Figure 6).

Telomere – From the point of emergence just after the posttorsal narrowing the proximal lobe of the telomere projects posteroproximad. It reaches above the primary coil of the solenomere at the height of the torsotope. The solenomere is wrapped into the telomere through an opening between the proximal lobe and the shallow lobe. The first half of the main stem of the telomere is closed in itself forming a sac-like structure. This sac opens at its posterior side resulting in an unfolding of the telomere into the two sheets. These sheets first curve anteriomesad and then proximad. Compared to the uncopulated telomere the sheets are less curled. On the inside of the telomere outgrowths emerge from the surface partially encasing the solenomere. Through this, the solenomere is held against the inner wall of the telomere and apically against the larger, posterior lamella of the telomere in particular (Figure 5B). Additionally, the telomere covers at least half of the bursal surface including the area where the solenomere tip is resting (Figure 6).

Vulvae (Figure 6)

The structure of the bursa is similar compared to the uncopulated state. However, the right and left bursae differ in their orientation. The furrow of the right bursa runs lateromediad while the furrow of the left bursa runs anteroposterad.

### MUSCLES OF THE GONOPODS AND VULVAE INVOLVED IN COPULATION

Various muscles associated with the gonopods and vulvae have been reconstructed in addition to the sclerotized structures to discuss possible muscle-mediated movements. They are depicted in Figure 6. A comprehensive list of all identified muscles including names and brief descriptions can be found in Table 1.

## DISCUSSION

### MATING BEHAVIOR

The mating behavior and copulatory sequence in *S. glabrus* is similar to most other Spirostreptida described so far (Telford and Dangerfield, 1993). As described by Telford and Dangerfield (1993), male Odontopygidae are very persistent in courtship, rapidly attaining a copula position and their copulation “appeared more physical” compared to many non-odontopygid Spirostreptida (Telford and Dangerfield, 1993: 3). This behavior may account for the short pre-copulatory and copulatory phase I. It has been described that sperm is translocated from the penes onto the gonopods during this phase (Telford and Dangerfield, 1993). Unfortunately, we were not able to observe the process of sperm translocation in *S. glabrus*. The alternating gonopod movements of *S. glabrus* during copulatory phase II, have been described for many other African Spirostreptida and Spirobolida (Barnett and Telford, 1996; Cooper and Telford, 2000; Telford and Dangerfield, 1993, 1996). It has been hypothesized that this behavior (1) serves as mechanical stimulation of the female to increase the chance of sperm acceptance, (2) removing the sperm of a precedent male and/or (3) to transfer the male’s sperm to the apodemic tube and were it is then somehow transferred into the spermathecae (Barnett, 1997). Using radioactively labelled males of *Doratogonus uncinatus* (Attems, 1914) (as *Alloporus uncinatus*) (Spirostreptidae) and *Zinophora diplodonta* (Attems, 1928) (as *Poratophilus diplodontus*) (Harpagophoridae) it has been shown that sperm transfer happens very early during copulation (Barnett and Telford, 1996; Barnett and Telford, 1994). Additionally, Barnett and Telford (1996) concluded based on dissections of copulated pairs that the telopodite of *D. uncinatus* is oriented correctly to facilitate sperm removal. Frederiksen and Enghoff (2015) reported the presence of sperm at the solenomere tip in *Raduliverpa serpentispina* Frederiksen & Enghoff, 2015 (Odontopygidae), proposing that this structure might facilitate sperm removal. The rhythmic gonopod movements of *S. glabrus* during copulatory phase II could facilitate both sperm transfer and sperm removal. Because gonopods of *S. glabrus* stay almost motionless during copulatory phase III, we argue that - similar to other Spirostreptida and Spirobolida - female stimulation, sperm mixture/removal and/or sperm transfer might take place during the previous copulatory phase II (Barnett, 1997; Cooper and Telford, 2000; Telford and Dangerfield, 1993). The long motionless copulatory phase III in *S. glabrus* fits well with former interpretations of such behavior as a form of mate guarding via prolonged copulation (Barnett, 1997; Parker, 1974; Telford and Dangerfield, 1996). The “unlocking” movement of male *S. glabrus* during copulatory-phase IV has also been described in other Spirostreptidae and has been hypothesized to be involved in the disconnection of gonopodal and vulval parts coupled during copulation. (Barnett, 1997). The post-copulatory vulval cleaning has also been reported from other Spirostreptida and could indicate post-copulatory female choice via selective sperm removal (Barnett and Telford, 1996; Telford and Dangerfield, 1990, 1996).

The bursae of *S. glabrus* correspond to the general description of bursae in Odontopygidae described by Barnett (1997) and we were not able to detect a significant difference between the uncopulated and copulated state. For this reason, we will focus on the gonopods solely in the following discussion.

### FUNCTIONAL MORPHOLOGY OF ODONTOPYGID GONOPODS

The functional morphology of helminthomorph gonopods in copula has been studied in Paradoxomatidae (Wojcieszek et al., 2012), Polydesmidae (Zahnle et al., 2020), Craspedosomatidae (Tadler, 1993) and Julidae (Tadler, 1996) using histology and/or μCT. In Spirostreptida (Spirostreptidae, Harpagophoridae and Odontopygidae), functional morphology of gonopods has been studied in copula by dissecting copulated pairs (Barnett, 1997; Demange, 1959; Kraus, 1966). The studies in Spirostreptida resulted in different hypotheses on the process of sperm translocation from the penes to the gonopods and sperm transfer from the gonopods to the bursae during copulation.

In Spirostreptida, non-motile sperm is translocated from the paired penes to the paired gonopods shortly prior to copulation. It is still unclear, whether sperm is translocated from the penes to the proximal base of the coxite (Gerhardt, 1933) or to its distal edge (Barnett, 1997; Brölemann, 1901; Demange, 1959; Kraus, 1966; Verhoeff, 1942). If translocated to the distal edge, sperm would have to travel proximad to the coxite base to enter the small (sperm-) channel of the telopodite. This might be due to gravity, if the male assumes an upright position. Additionally, movements of the basomere part of the telopodite could act as a plunger pumping the sperm distad through the metaplical fold (Kraus, 1966; Verhoeff, 1942). The same movement could also be involved in pumping sperm up the sperm channel (Kraus, 1966; Verhoeff, 1942). Alternatively, in Spirostreptidae, the solenomere could pick up sperm directly from the distal edge of the coxite via gonopodal movements during the retraction and release cycle (Barnett, 1997; Barnett and Telford, 1996). Barnett and Telford (1996) also suggest that this is not possible in Harpagophoridae and Odontopygidae, where the solenomere does not make contact to the distal edge of the coxite during gonopodal movements. We were not able to observe sperm translocation in *S. glabrus*. However, based on our morphological interpretations we favour the hypothesis that sperm is translocated to the distal edge of the coxite and then pumped down the metaplical fold and up the sperm channel all the way to the solenomere tip. (1) As mentioned by Kraus (1966) the distal edge of the coxite is more or less funnel-shaped and exactly fits the form of the penis supporting an efficient translocation process to this gonopodal region. (2) The lateral apodeme-telopodite muscle (gm3) may be able to drag the telopodite proximad against the hydrostatic pressure of the gonopodal pillow. This would result in the basomere perfectly acting as the plunger pumping sperm down the metaplical fold as proposed by Verhoeff (1942) and Kraus (1966).

Subsequently to sperm translocation, the sperm has to be transferred from the gonopods to the bursae. It is accepted that, prior to sperm translocation and copulation, gonopods are everted hydrostatically from the gonopodal cavity by increasing haemolymph pressure in the gonopodal pillow (Barnett, 1997; Brölemann, 1901; Demange, 1959; Gerhardt, 1933; Kraus, 1966; Verhoeff, 1942). Verhoeff (1926-32: 184) suggested an additional function of the gonopodal pillow (“Blähpolster” *sensu* Verhoeff). However, the mechanism of copulation in general, the gonopod erection and the sperm transfer compared to sperm translocation is still under discussion. In Spirostreptida, basomere retraction in the everted gonopod leads to a de-spiralization of the torsotope. This results in a multi-step spinning movement of the solenomere and its subsequent erection (Barnett and Telford, 1996; Demange, 1959; Kraus, 1966). While the subsequent de-spiralization of the torsotope has been reported for all Spirostreptida observed, an erection of the solenomere has been reported for Spirostreptidae and Harpagophoridae but not for Odontopygidae by Barnett and Telford (1997) (for Spirostreptidae see also Brölemann, 1901; Verhoeff, 1942 and Demange, 1959). Kraus (1966) however mentioned that the solenomere of Odontopygidae observed by him also erects during copulation. These different reports of solenomere erection in Odontopygidae could be explained by the techniques used by Barnett and Telford. These authors manipulated the gonopodal muscles mechanically to induce gonopod movements. Possible hydrostatic aspects of the system have not been tested by these authors. Our 3D reconstruction of the copulating *S. glabrus* pair clearly shows an erected solenomere agreeing with Kraus (1966) and contradicting the descriptions by Telford and Barnett (1996) and Barnett (1997). This indicates a conserved gross mechanism underlying gonopod function within Spirostreptida.

Different forces have been proposed to underlay torsotope de-spiralization and solenomere erection. Kraus (1966) suggested that muscle-mediated basomere retraction induces stress on the remaining parts of the telopodite. The elastic properties of the heterogeneously sclerotized torsotope (“Aufweichzone” *sensu* Kraus, 1966) transform this stress into the observed de-spiralization movement and the subsequent solenomere erection. Demange (1959) additionally suggested the prostatic gland (“glandes prostatiques” *sensu* Brölemann, 1921; “Basaldrüse” *sensu* Verhoeff, 1942) - the large channel running from the basomere, through the torsotope into the solenomere - could serve as a pressure reservoir contributing to solenomere erection. This gland has been proposed to add fluid secretion to the sperm and either flush it up the sperm channel or diluting it so that it can be pumped up more easily through the respective channel (Brölemann, 1901; Kraus, 1966; Verhoeff, 1942). In Chordeumatida, Haacker (1971) and Verhoeff (1926-32: 185) reported that a similar gland produces a secretion necessary for spermatophore production. Yet, the presence of a spermatophore has been rejected for this group by Tadler (1993). There are at least three possibilities of the role of the large channel in the telopodite of Odontopygidae based on the descriptions by previous authors and our morphological observations. (1) It is a gland producing sperm secretion supplements not involved in mechanical gonopod function. (2) It has no gland function but rather acts as a pressure reservoir only. (3) It has a double function, serving as a gland producing sperm supplements and emptying before copulation. After emptying it offers the space to serve as a pressure reservoir. The glandular nature of the large channel has to be examined histologically and ultrastructurally, and a possible hydrostatic role involved in telopodite de-spiralization and/or solenomere erection has to be tested experimentally to solve this question. However, the observation that the chamber of the large channel near the torsotope increases in volume in the erected compared to the non-erected state supports the function as such a hydrostatic reservoir.

Furthermore, the function of the solenomere tip in Spirostreptida is also under discussion. This structure has been proposed to serve in sperm transfer (Kraus, 1966) as well as sperm removal or mixture (Barnett and Telford, 1996). Evidence for its function in sperm transfer is based on the presence of a small channel (sperm channel) running from the telopodite basomere to the solenomere tip. During copulation, the solenomere tip is located above the apodemic tube of the female bursa, allowing the sperm to be transferred in the proximity of the spermathecae. However, this position above the apodemic tube, the diverse denticulation of the solenomere tip including different spines and the presence of spermatozoa on its surface have been used as evidence for its role in sperm removal or mixture (Barnett and Telford, 1996; Frederiksen and Enghoff, 2015). Alternatively, it has been proposed for Spirostreptidae that sperm is transferred to the bursa via the distal edge of the coxite. Afterwards movements of the now erected solenomere relocate the sperm closer to the spermathecae of the female bursae (Barnett et al., 1995). We argue that in *S. glabrus* it is unlikely that sperm is transferred to the bursa via the distal edge of the coxite. Such a process would involve the intromission of the everted but non-erected gonopods into the vulval sac. Subsequently, they would have to be completely pulled out again to erect the solenomere due to the limited space inside the vulval sac. Thereafter, another round of gonopod intromission would continue with the erected solenomere to relocate or mix the previously placed sperm.

While most functional discussions focus on the coxite, the large telopodite channel, the torsotope and the solenomere, possible functions of the paracoxite and the telomere have not been discussed to our best knowledge. We therefore incorporate own functional hypotheses of these structures to propose an elaborate model of gonopod function in Odontopygidae based on the model of Barnett and Telford (1996) (Figure 7).

**Figure 7.**
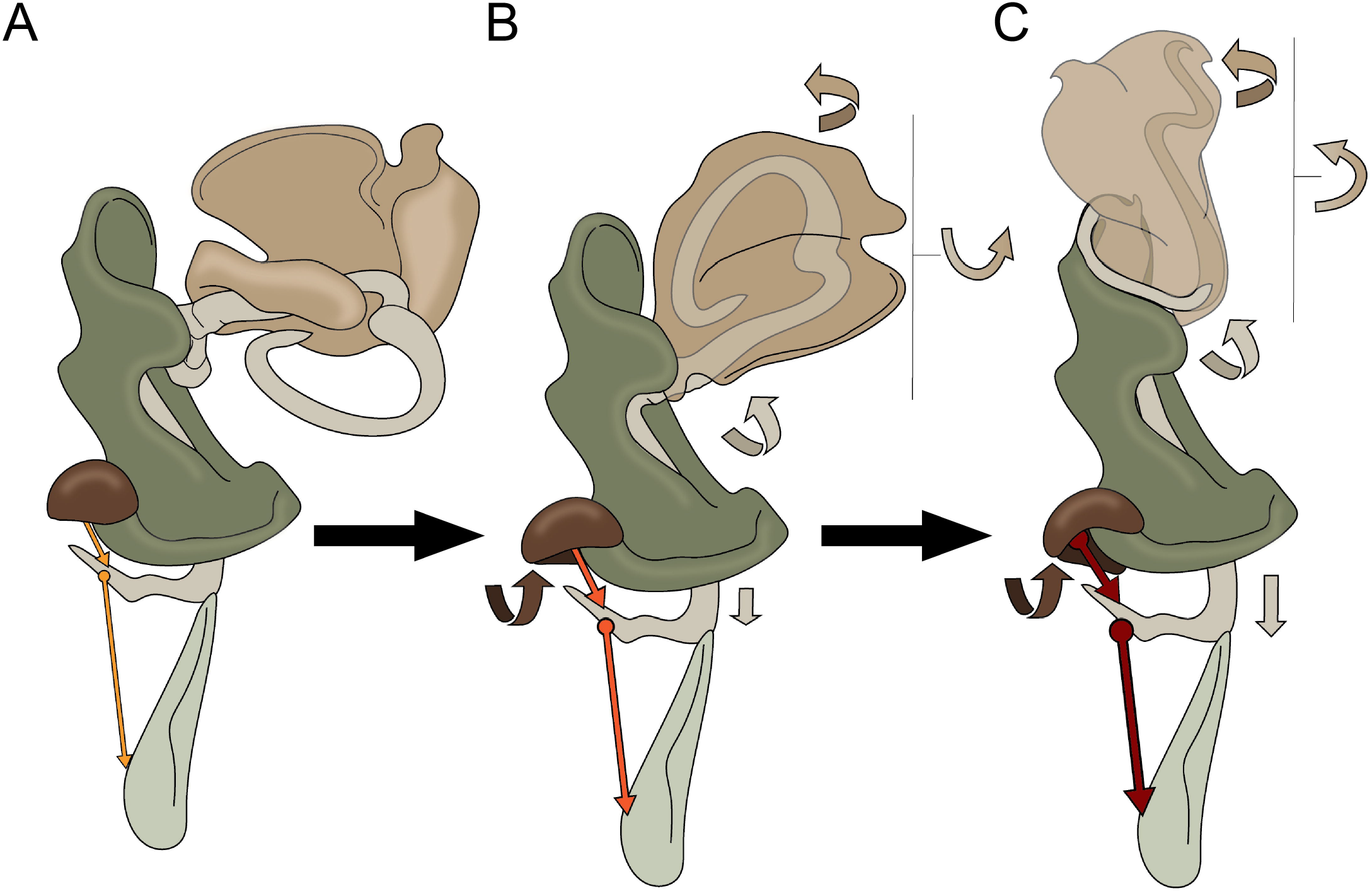
Schematic drawing of the gonopod of *Spinotarsus* in a resting/uncopulated (A), intermediate (B) and copulated phase (C). Small arrows indicate the movements of parts of the corresponding color. The orange-to-red shaded arrows indicate contraction states of the gm3 and gm4. Details are given in the text.

### AN ELABORATED FUNCTIONAL MODEL OF THE COPULATORY MECHANISM OF ODONTOPYGIDAE

At the beginning of copulation, the gonopods are situated in a resting position inside the gonopodal sac (Figure 7A). To initiate copulation, hemolymph is pumped into the gonopodal pillow pushing the gonopods out of the gonopodal sac. Subsequently, the lateral apodeme-telopodite muscle (gm3) contracts, displacing the basomere distad, pulling it down through the metaplical fold. This induces stress to the sclerotized wall of the torsotope. The interactions of the stressed, strongly sclerotized areas of the torsotope and the less stressed, weakly sclerotized areas (“Aufweichzone” *sensu* Kraus, 1966) results in torsotope de-spiralization and solenomere erection (Figure 7B). Provided that the large channel acts as a pressure reservoir, hemolymph is pushed simultaneously into the large channel of the telopodite. This leads to an increased volume and induced pressure on the torsotope walls that could complement the tension induced by gm3 contraction, and the proximal displacement of the basomere. The erected solenomere is wedged beneath the cucullus increasing the resistance of the solenomere erection against pressure fluctuations within the gonopodal pillow (Figure 7C). Additional fixation of the erected solenomere is facilitated by the contraction of the paracoxite-telopodite muscle (gm4) rotating the paracoxite around the basal lateral coxal process. In the rotated position, the paracoxite and the gm4 block the space between the telopodite arm and the coxa, preventing the basomere from sliding back into the metaplical fold. During intromission into the vulval sac and due to the limited space therein, the solenomere is enwrapped by the telomere lamellae. At this point, the telomere acts as a guiding rail protecting the delicate solenomere. Additionally, the weakly sclerotized telomere might ease keeping up the copula position by pressing against the inner vulval sac due to material-mediated elastic properties and thereby wedging and fixing the gonopods inside the female. The complete movements bring the tip of the solenomere close to the apodemic tube of the bursa. The combination of gm3 contraction and slight pressure fluctuation of the gonopodal pillow may easily allow both, pumping of sperm up the sperm channel and mixing or scraping movements of the solenomere tip as suggested by several authors (Barnett and Telford, 1996; Demange, 1959; Frederiksen and Enghoff, 2015; Kraus, 1966). To end copulation, pressure in the gonopodal pillow is lowered. Contraction of the distal apodeme-coxa muscle (gm1) and the apodeme tergite muscle (gm2) facilitate withdrawal of the gonopods out of the vulval sac. Decreasing pressure and gm1 contraction also result in the restoration of the solenomere resting configuration that might correspond to the “unlocking” movement observed during copulatory phase IV. Lastly, after facultative cleaning of the gonopods by the male’s mouthparts, the ventral t6-gonopodal sac muscle (gm5) and the dorsal t6-gonopodal sac muscle (gm6) open the gonopodal sac and the gonopods are stored there by further contraction of the gm1 and gm2.

### MECHANISMS UNDERLYING GONOPOD EVOLUTION

Hypotheses trying to explain the evolution of gonopod disparity aim to identify the underlying selective mechanisms: lock-and-key mechanism (Wojcieszek et al., 2012; Zahnle et al., 2020), pleiotropy (Kraus, 1966), sperm competition (Barnett and Telford, 1996) and/or female choice (Tadler, 1993). Despite all the differences between gonopods of Odontopygidae, their basic form might be constrained primarily by basic functional demands facilitating sperm translocation and transfer. The cucullus serves as a funnel receiving the sperm from the penis; the metaplical fold, the basomere and the gm4 act as a plunger moving the sperm down the coxa and then up the telopodite within the sperm channel; the gm3 and the torsotope allowing a conformational change to store the gonopods inside the gonopodal sac. Especially the last function is crucial to protect the delicate gonopods of species burrowing through the substrate. We found no evidence for a close morphological correspondence of the copulated genitals in *S. glabrus* as reported for Paradoxomatidae (with a morphologically simple vulva; Wojcieszek et al., 2012) and Polydesmidae (with a morphologically complex vulva; Zahnle et al., 2012). An experimental study on separated populations of the paradoxomatid *Antichiropus variabilis* (Attems, 1911) comparing genetic structure, gonopod morphometry and paternity success after mating of two of the most divergent populations found good arguments for the action of stabilizing, hinting to a possible lock-and-key mechanism in this species (Wojcieszek and Simmons, 2012). However, a geometric characterization of the bursae has not been carried out in this study. Therefore it might be possible that the decreased paternity success in mating experiments of two distant *A. variabilis* populations is due to increased genetic incompatibility rather than a mechanical one. Much more studies such as the pioneering ones by Wojcieszek and Simmons (Wojcieszek and Simmons, 2011, 2012, 2013) combined with detailed data on the functional morphology has to be carried out to further test the presence of a lock-and-key mechanism underlying copulatory organ evolution in different helminthomorph groups. However, from a morphological point of view, the relatively simple structure of the odontopygid bursa makes it unlikely that genital evolution within Odontopygidae (and maybe Spirostreptida in general) was driven by selection for a lock-and-key mechanism. Additionally, on the basis of theoretical considerations and morphological data, it is problematic to distinguish a lock-and-key mechanism from female choice for best fit. The result of both selective mechanisms would appear more or less similar – copulatory organs that somehow fit each other. We therefore argue that female choice and sperm competition are the most important mechanisms underlying gonopod evolution of the Odontopygidae and maybe of Spirostreptida in general. A first indicator for female choice is the observed repelling behavior of the female and the rigorous mounting behavior of the male trying to attain the mating position. Additionally, the gonopods of *S. glabrus* exhibit many features involved in stabilization and maintenance of the copulatory position against female repelling behavior - the paracoxite and the gm4 block a backwards slide of the telopodite into its resting position; the wedged edge of the cucullus fixes the erected solenomere and the telomere protects the delicate solenomere and anchors the gonopods inside the vulval sac. While there is no close correspondence between these structures and the female bursa as demanded for a lock-and-key mechanism, they are involved in maintaining the copula against female disturbance as proposed by female choice via male-female conflict (Cordero and Eberhard, 2003). An additional mechanism influencing genital morphology is sperm competition (Parker, 1970). The presence of sperm removal in some Spirostreptida via the solenomere has been proposed by different authors (Barnett and Telford, 1996; Frederiksen and Enghoff, 2015). It has been shown in some Spirostreptida that (1) the sperm of a precedent male can be partially removed by a following male via sperm flushing and/or, superficially, by scraping parts of the bursal apodemic tube with the solenomere tip (Barnett, 1997). This is in accordance with our observation of the position of the solenomere tip during copulation allowing both, a scraping movement and sperm flushing via the small solenomere (sperm) channel. Additionally, females might be able to selectively discard “unwanted” sperm (Barnett et al., 1995). However, an innervation of the bursa, which would be crucial for such an action, has not been shown yet.

## ACKNOWLEDGEMENTS

We would like to thank Lars Möckel and Hendrik Müller for help collecting the specimens investigated in this study. We are grateful to Stephan Scholz for the preparation of μCT-scans and to Christoph Höpel for help with the CLSM. We thank Christian Wirkner for fruitful discussions and Stefan Richter for critical reading of the manuscript.

## FIGURE CAPTURES

**Table 1.** Musculature associated with the copulatory organs of male and female *S. glabrus*. T, trunk segment (followed by a number, e.g., t6 means trunk segment number 6).

**Supplementary Material 1**. Time values of mating phases of *S. glabrus*. “–”, time not recorded; min., minutes (measured in 5 minute intervals); sec., seconds.

**Supplementary Material 2**. Video recording of mating in *Spinotarsus glabrus*.

## REFERENCES

Attems, C., 1914. Afrikanische Spirostreptiden nebst Überblick über die Spirostreptiden orbis terrarum. Zoologica (Stuttgart) 25 (65-66), 1–235.

Attems, C., 1950. Über Spirostreptiden (Diplopoda). Annalen des Naturhistorischen Museums in Wien 57, 179–257.

Barnett, M., Telford, S.R., 1994. The timing of insemination and its implications for sperm competition in a millipede with prolonged copulation. Animal Behaviour 48 (2), 482–484.

Barnett, M., Telford, S-R., Tibbles, B.J., 1995. Female mediation of sperm competition in the millipede Alloporus uncinatus (Diplopoda: Spirostreptidae). Behavioral Ecology and Sociobiology 36 (6), 413–419.

Barnett, M., Telford, S.R., 1996. Sperm competition and the evolution of millipede genitalia. -Mémoires du Muséum national d’histoire naturelle, N. S. 169, 331–339.

Barnett, M., 1997. Sex in southern african spirostreptida millipedes: mechanisms of sperm competition and cryptic female choice. Thesis. University of Cape Town. 177pp.

Bock, W.J., von Wahlert, G., 1965. Adaptation and the form-function complex. Evolution, 269–299.

Brölemann, H.W., 1901. Materiali per la conoscenza della Fauna Eritrea raccolti dal Dott. P. Magretti. Myriapodes. Bollettino della Società entomologica Italiana 33 (1): 26–35.

Cooper, M.I., Telford, S.R., 2000. Copulatory sequences and sexual struggles in millipedes. Journal of Insect Behavior 13 (2): 217–230.

Cordero, C., Eberhard, W.G., 2003. Female choice of sexually antagonistic male adaptations: a critical review of some current research. Journal of Evolutionary Biology 16, 1–6.

Demange, J.-M., 1959. L’accouplement chez Graphidostreptus tumuliporus (Karsch) avec quelques remarques sur la morphologie des gonopodes et leur fonctionnement. Bulletin de la Société entomologique de France 64, 198–207.

Eberhard, W.G., 1985. Sexual selection and animal genitalia. Harvard University Press. 244pp.

Eberhard, W.G., 1993. Evaluating models of sexual selection: genitalia as a test case. The American Naturalist 142, 564–571.

Frederiksen, S.B., Enghoff, H., 2012. East African odontopygid millipedes 1: Five new species of Xystopyge (Attems, 1909) and a proposal for a new gonopod terminology (Diplopoda; Spirostreptida; Odontopygidae). Zootaxa 3485: 69–82.

Frederiksen, S.B., Enghoff, H., 2015. East African odontopygid millipedes 4: A restricted redefinition of the genus Rhamphidarpoides Kraus, 1960, a related new genus, five new species, and notes on solenomere function (Diplopoda; Spirostreptida; Odontopygidae). Zootaxa 3926, 541–560.

Gerhardt, U., 1939. Zur Funktion der Gonopoden bei Graphidostreptus gigas (Peters) (Diplop. Julif.). Mitteilungen aus dem Zoologischen Museum in Berlin 19, 430–439.

Kraus, O., 1966. Phylogenie, Chorologie und Systematik der Odontopygoideen (Diplopoda, Spirostreptomorpha). Abhandlungen der Senckenbergischen naturforschenden Gesellschaft 512, 1–143.

Kraus, O., 1968. Isolationsmechanismen und Genitalstrukturen bei wirbellosen Tieren. Zoologischer Anzeiger 181 (1-2), 22–38.

Minelli, A., 2015. Treatise on Zoology - Anatomy, Taxonomy, Biology. The Myriapoda, Volume 2 – Diplopoda. Brill, Leiden. 482pp.

Parker, G.A., 1970. Sperm competition and its evolutionary consequences in the insects. Biological Reviews 45, 525–567.

Parker, G.A., 1974. Courtship persistence and female-guarding as male time investment strategies. Behaviour 48, 157–183.

Schindelin, J., Arganda-Carreras, I., Frise, E., Kaynig, V., Longair, M., Pietzsch, T., Preibisch, S., Rueden, C., Saalfeld, S., Schmid, B., 2012. Fiji: an open-source platform for biological-image analysis. Nature Methods 9, 676–682.

Tadler, A., 1993. Genitalia fitting, mating behaviour and possible hybridization in millipedes of the genus Craspedosoma (Diplopoda, Chordeumatida, Craspedosomatidae). Acta Zoologica 74 (3), 215–225.

Tadler, A., 1996. Functional morphology of genitalia of four species of julidan millipedes (Diplopoda: Nemasomatidae; Julidae). Zoological Journal of the Linnean Society 118 (1), 83–97.

Telford, S.R., Dangerfield, J.M., 1990. Sex in millipedes: laboratory studies on sexual selection. Journal of Biological Education 24, 233–238.

Telford, S.R., Dangerfield, J.M., 1993. Mating behaviour and mate choice experiments in some tropical millipedes (Diplopoda: Spirostreptidae). South African Journal of Zoology 28 (3), 155–160.

Telford, S.R., Dangerfield, J.M., 1996. Sexual selection in savanna millipedes: products, patterns and processes. Mémoires du Muséum national d’histoire naturelle, N. S. 169, 565–576.

Verhoeff, K.W., 1926-1932. Gliederfüßler: Arthropoda, II. Abteilung: Myriapoda. 2. Buch: Diplopoda. Klassen und Ordnungen des Tierreichs 5 (2), 1–2084.

Verhoeff, K.W., 1942. Myriapoden der Insel Fernando Po und über den Ankerapparat und die Spermaleitung der Spirostreptoideen. XVI. Beitrag zu den wissenschaftlichen Ergebnissen der Forschungsreise H. Eidmann nach Spanisch-Guinea, 1939/40. Zeitschrift für Morphologie und Ökologie der Tiere 39 (1), 76–97.

Wilson, H.M., 2002. Muscular anatomy of the millipede Phyllogonostreptus nigrolabiatus (Diplopoda: Spirostreptida) and its bearing on the millipede “thorax”. Journal of Morphology 251 (3), 256–275.

Wojcieszek, J.M., Austin, P., Harvey, M.S., Simmons, L.W., 2012. Micro-CT scanning provides insight into the functional morphology of millipede genitalia. Journal of zoology 287 (2), 91–95.

Wojcieszek, J.M., Simmons, L.W., 2011. Male genital morphology influences paternity success in the millipede Antichiropus variabilis. Behavioral Ecology and Sociobiology 65, 1843–1856.

Wojcieszek, J.M., Simmons, L.W., 2012. Evidence for stabilizing selection and slow divergent evolution of male genitalia in a millipede (Antichiropus variabilis). Evolution 66, 1138–1153.

Wojcieszek, J.M., Simmons, L.W., 2013. Divergence in genital morphology may contribute to mechanical reproductive isolation in a millipede. Ecology and Evolution 3, 334–343.

Zahnle, X.J., Sierwald, P., Ware, S., Bond, J.E., 2020. Genital morphology and the mechanics of copulation in the millipede genus Pseudopolydesmus (Diplopoda: Polydesmida: Polydesmidae). Arthropod Structure and Development 54, 100913.

